# Substrate stiffness and viscoelasticity influence fibroblast senescence

**DOI:** 10.1101/2025.08.22.671571

**Authors:** Mackenzie L. Skelton, Tanvi Bhat, Ethan Yu, Afia Ofori, Steven R. Caliari

## Abstract

Senescent cell accumulation has been implicated in aging and fibrotic disease, which are both characterized by increased tissue stiffness. However, the direct connection between tissue mechanics and senescence induction remains disputed in the literature. Thus, this work investigates the influence of hydrogel stiffness and viscoelasticity in promoting fibroblast senescence both in combination with genotoxic stress and independently. We show that while lung fibroblast YAP signaling declines with senescence induction, senescent fibroblasts maintain their mechanosensing capabilities with increased YAP nuclear localization on higher stiffness hydrogels. Most notably, we find a unique role for hydrogel viscoelasticity in senescence induction with soft (2 kPa) viscoelastic substrates promoting both the onset and amplification of senescence, even in the absence of genotoxic stress. These changes are not associated with a decline in YAP activity, but instead with a decline in nuclear DAPI intensity, suggesting a role of nuclear organization in driving this phenotype. Overall, this work highlights the influence of mechanics on the induction of senescence and supports the key role of viscoelasticity.

## Introduction

Cellular senescence is a hallmark of aging tissue and is characterized by persistent cell cycle arrest, cell and nuclear enlargement, and a pro-inflammatory secretory profile^1,2^. The accumulation of senescent cells has been implicated in driving many devasting, age-related diseases such as idiopathic pulmonary fibrosis (IPF)^3,4^. In parallel, there are various compositional and mechanical changes to the extracellular matrix (ECM) with age and disease, such as elevated collagen content leading to increased stiffness^5–7^. However, whether these changing mechanics contribute to the senescent cell burden is not known.

A decline in Yes-associated protein 1 (YAP) activity, a master regulator of mechanotransduction, has been well documented with senescence^8–10^. YAP signaling has been reported as the gatekeeper of cellular senescence in stromal cells through the preservation of nuclear envelope integrity^8^. Additionally, others have reported that changes in YAP activity are responsible for the morphological changes and secretory profile of senescent cells^9^. However, these studies relied on genetic manipulation of YAP rather than mechanically-mediated changes in YAP activity. Thus the interaction between stiffness-mediated YAP activation and senescence-induced YAP decline is yet to be elucidated.

Previous studies investigating the role of elastic modulus, or stiffness, on senescence phenotype showed mixed results. Various studies reported evidence of substrates with stiffnesses of 15 kPa and above enhanced the senescent phenotype of stromal cells^11–13^. Further, one study showed evidence that hydrogels with a stiffness of 16 kPa were sufficient on their own to induce senescence in vascular smooth muscle cells^14^. Conversely, other studies reported that soft substrates, ranging from 0.5 kPa to 6 kPa, either enhance or induce the senescent phenotype in stromal cells^15,16^. Yet, even within these studies there are variable findings. One study found that passage 2 primary rat cardiac fibroblasts show increased levels of senescence on 6 kPa hydrogels, while passage 5 cells show equally high levels of senescence on 6 or 40 kPa substrates, and freshly isolated cells show no senescence on either hydrogel^16^. Another study showed that transferring cells from 0.5 kPa substrates to 25 kPa substrates elevated levels of p16 and p21, two canonical markers of cellular senescence, arguing that these biomarkers are mechanically regulated^15^. Taken together, these studies highlight the need for clarity as to whether YAP-mediated mechanotransduction regulates senescence in the context of substrates that mimic tissue mechanics.

In addition to stiffness, viscoelasticity has emerged as not only a key mechanical characteristic of all human tissues, but also an important regulator of cell fate^17–19^. Viscoelasticity describes the time-dependent ability of a material to display properties of both solids and liquids, such as stress relaxation^17^. A recent report showed elegantly that YAP nuclear translocation was more sensitive to energy dissipation than to substrate rigidity^20^, emphasizing the importance of this mechanical component on cell behavior and senescent phenotype. One recent study claimed viscoelasticity triggered cellular clustering that could be protective against senescence induction^21^. However, given that cell-cell contact reduces YAP nuclear localization independent of mechanics^22^, there is a need to decipher cellular responses to viscoelastic mechanics alone.

In this work, we systematically investigated the influence of substrate stiffness and viscoelasticity on the induction and maintenance of the senescent phenotype of primary human lung IMR90 fibroblasts. We utilized a high-throughput hydrogel platform with well-defined and tunable hyaluronic acid-based hydrogels to modulate substrate stiffness and viscoelasticity independently within a single 96-well plate. We leveraged genotoxic stress to induce cellular senescence through treatment with bleomycin and confirmed the development of a robust senescent phenotype with a concurrent reduction in YAP activity. Viscoelastic substrates were shown to play a unique role in senescence induction both independently and synergistically in the presence of genotoxic stress. Overall, this work provides novel insight into the role of substrate mechanics in inducing fibroblast senescence.

## Results

### Bleomycin treatment induces a robust senescence phenotype in fibroblasts

IMR90 fibroblasts were exposed to genotoxic stress via bleomycin to induce senescence. A variety of dosing regimens were tested to identify the most effective approach to generate high levels of senescent cells. We found that the timing of bleomycin treatment had a greater impact on senescent levels than the dose, with dose-dependent effects seen only at the 24 hour time point, but not at 2 or 72 hours (Fig. S1). Thus, a low dose of 20 µg/mL bleomycin was used, and cells were treated for 72 hours with an additional 24 hour culture to minimize cell death and maximize the number of senescent cells.

Bleomycin-treated cells showed high levels of the canonical senescent cell marker β-galactosidase (β-gal), with greater than 80% of cells staining positive, compared to less than 5% in the untreated control (Fig. 1a). These cells showed significant increases in overall spreading and nuclear area (Fig. 1b), a common characteristic of senescent cells^9,23^. Additionally, cells treated with bleomycin displayed a marked decline in proliferation as measured by Ki67 expression (Fig. 1c). Bleomycin treatment increased the expression of cell cycle regulator p21 (Fig. 1d), and common senescence-associated secretory phenotype (SASP) components were upregulated at the RNA level (Fig. 1e). These data provide ample confidence in this treatment regime for the thorough induction of senescence.

**Figure 1:**
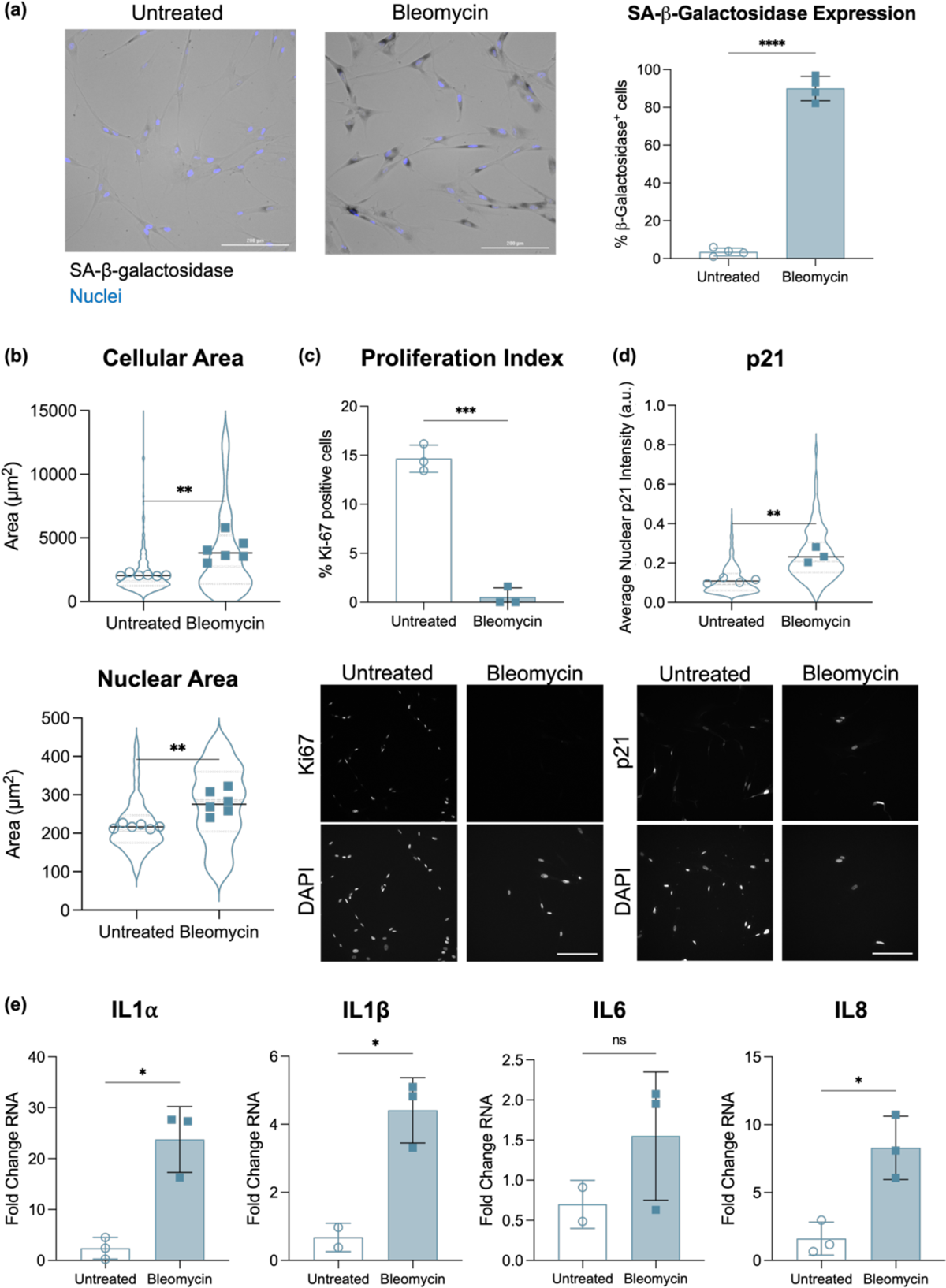
Bleomycin induces senescence in IMR90 fibroblasts. (a) Representative images of fibroblasts either untreated or treated with bleomycin and stained with senescence-associated (SA)-β-gal and DAPI to stain nuclei. Quantification of the number of cells that stained positive for β-gal is shown on the right, indicating there was significantly greater expression of β-gal in bleomycin-treated cells. 143 and 332 cells were counted cells for the bleomycin-treated and untreated groups respectively with N = 4 biological replicates each. (b) Cell and nuclear area both significantly increased with bleomycin treatment. 200 and 766 cells were analyzed for the bleomycin-treated and untreated groups respectively, with N = 6 biological replicates each. (c) Cells showed a significant decrease in proliferation when treated with bleomycin as measured by Ki67 expression. 389 and 876 cells were analyzed for the bleomycin-treated and untreated groups respectively, with N = 3 biological replicates each. Representative images shown below. (d) p21 expression significantly increased in cells treated with bleomycin. 163 and 464 cells were analyzed for the bleomycin-treated and untreated groups respectively, with N = 3-4 biological replicates. Representative images shown below. (e) RT-qPCR showed significant increases in IL1α, IL1β, and IL8 expression. N = 2-3 replicates. Scale bars: 200 µm. All statistical tests are Welch’s t-tests. *p<0.0332, ** p<0.0021, ***p< 0.0002, ****p< 0.0001.

### YAP activity declines with bleomycin treatment

We next sought to confirm whether YAP signaling declines with bleomycin treatment, as others have shown with the onset of senescence^8–10^. As expected, YAP nuclear localization shows a significant decline with bleomycin treatment as seen in example images (Fig. 2a) and quantification of average nuclear to cytoplasmic YAP intensity ratio (Fig. 2b). Further, a decrease in YAP nuclear localization is correlated with an increase in p21 expression (Fig. 2c, p = 0.035), emphasizing the concurrent decline in YAP activity with increased expression of senescence markers. We also noted a significant decrease in average nuclear DAPI intensity upon bleomycin treatment (Fig. 2d), which may be correlated with the hypothesis that declining YAP nuclear localization is caused by a disruption in nuclear membrane integrity^8^. Additionally, a decline in nuclear DAPI intensity has been shown to indicate a more open chromatin structure^24^. This decline in nuclear DAPI signal is also strongly correlated with the decline in nuclear YAP (Fig. 2e, p = 0.016). Taken together, these data support that this model system aligns with previous observations regarding a decline in active YAP signaling with senescence.

**Figure 2:**
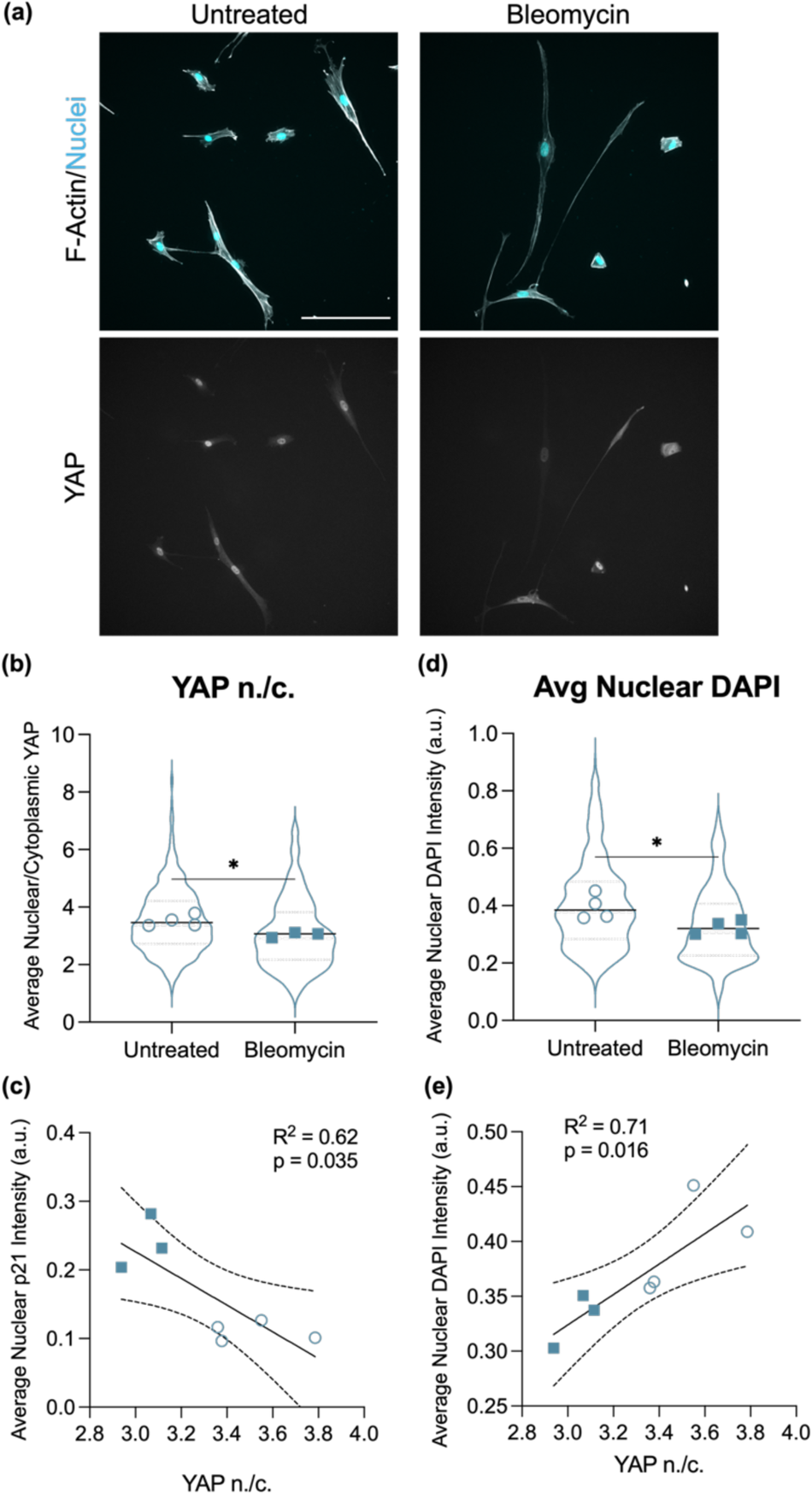
YAP activity declines with bleomycin treatment. (a) Representative images of fibroblasts either untreated or treated with 20 µg/mL bleomycin. Scale bar: 200 µm. (b) Quantification of average nuclear (n.) to average cytoplasmic (c.) YAP ratio showed a decrease in nuclear YAP in cells treated with bleomycin. (c) YAP n./c. trended inversely with average nuclear p21 intensity. (d) Quantification of average nuclear DAPI intensity showed a decline in signal with bleomycin treatment. (e) Average nuclear DAPI intensity correlated strongly with YAP n./c. Statistics for (b) and (d) are student’s t-tests where *p<0.0332. N = 3-4 replicates per group with 163-464 cells per group quantified.

### Senescent fibroblasts retain mechanosensing capabilities

As YAP is a master mediator of mechanotransduction, we next investigated whether senescent fibroblasts with blunted YAP activity also maintain functional mechanosensing capabilities. Fibroblasts were cultured in flasks either dosed with 20 µg/mL bleomycin or left untreated. After 3 days, both treated and untreated flasks had media replaced to normal culture media, then 1 day later cells were passaged onto hydrogels of varying mechanics. Hydrogels were fabricated with stiffnesses ranging from 2 kPa to 40 kPa through an increase in polymer concentration and/or crosslinking density, and 2 kPa viscoelastic hydrogels were fabricated through the introduction of physical guest-host crosslinks to the polymer network^25–27^. Cells were cultured on hydrogels for 24 hours before fixing and staining (Fig. 3a). Cells were stained to visualize F-actin to assess cell morphology and F-actin fiber organization, nuclei, and YAP (Fig. 3b). Cells that were untreated maintained expected behavior with an increase in YAP nuclear localization with increasing stiffness (Fig. 3c), more elongated morphology (lower form factor, Fig. 3d), and greater F-actin organization (lower actin uniformity, Fig. 3e). Cells treated with bleomycin also maintained these stiffness-dependent trends, although there were significant differences between the untreated and bleomycin-treated cells seeded on the same stiffnesses (Fig. S2). Specifically, YAP activity declined significantly with bleomycin treatment for cells seeded on 10 and 40 kPa substrates, but these differences were less pronounced on softer hydrogels (Fig. S2a). Differences in YAP activity on 2 kPa viscoelastic and elastic hydrogels were not statistically different. Further, bleomycin-treated cells trended toward assuming a myofibroblast-like morphology on all hydrogel groups, with significantly greater cellular elongation and F-actin organization (Fig. S2b-c).

**Figure 3:**
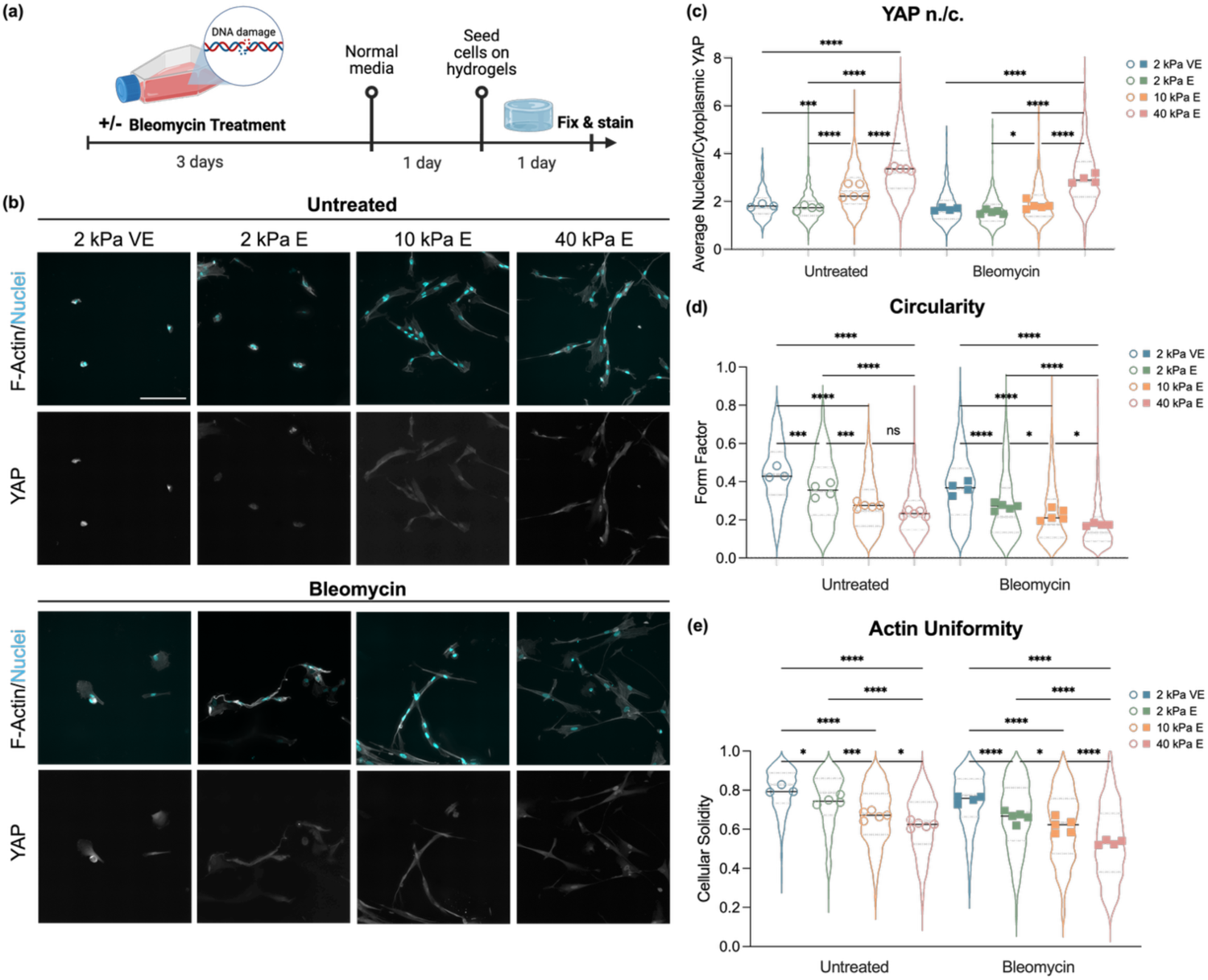
Senescent fibroblasts retain mechanosensing capabilities. (a) Schematic overview of experimental workflow. (b) Representative images of F-actin and DAPI overlay or YAP staining of fibroblasts on 2 kPa viscoelastic (VE), 2 kPa elastic (E), 10 kPa E, and 40 kPa E hydrogels, either untreated or treated with bleomycin. Scale bar: 200 µm. In both untreated and bleomycin-treated cells (c) YAP n./c. increased as a function of stiffness, (d) cellular morphology became less circular, and (e) actin uniformity decreased, indicating an increase in stress fiber formation. Two-way ANOVA tests were performed with Tukey’s multiple comparison test. N = 3-5 hydrogels per group with 367-1501 cells per group quantified. *p<0.0332, ***p<0.0002, ****p<0.0001.

### Fibroblasts treated with bleomycin retain senescent characteristics when plated on hydrogels

We next sought to confirm our assumption that fibroblasts treated with bleomycin retained their senescent phenotype when plated on hydrogels. Fibroblast β-gal expression was assessed 24 hours after plating on hydrogels. We additionally measured cell and nuclear area, as well as average DAPI intensity. As expected, cells treated with bleomycin maintained high levels of β-gal activity once seeded on hydrogels (Fig. 4a-b), indicating that the exposure to softer hydrogel substrates does not reverse an already established senescence phenotype. This is also reflected in general increases in cell and nuclear areas (Fig. 4c-d) as well as a consistent decrease in average nuclear DAPI (Fig. 4e). Interestingly, cells that were not exposed to genotoxic stress showed non-negligible levels of β-gal staining when seeded on 2 kPa viscoelastic hydrogels (Fig. 4b). This trend toward a senescent phenotype was corroborated by the lack of a significant change in nuclear area between bleomycin-treated and untreated cells on 2 kPa viscoelastic substrates (Fig. 4d).

**Figure 4:**
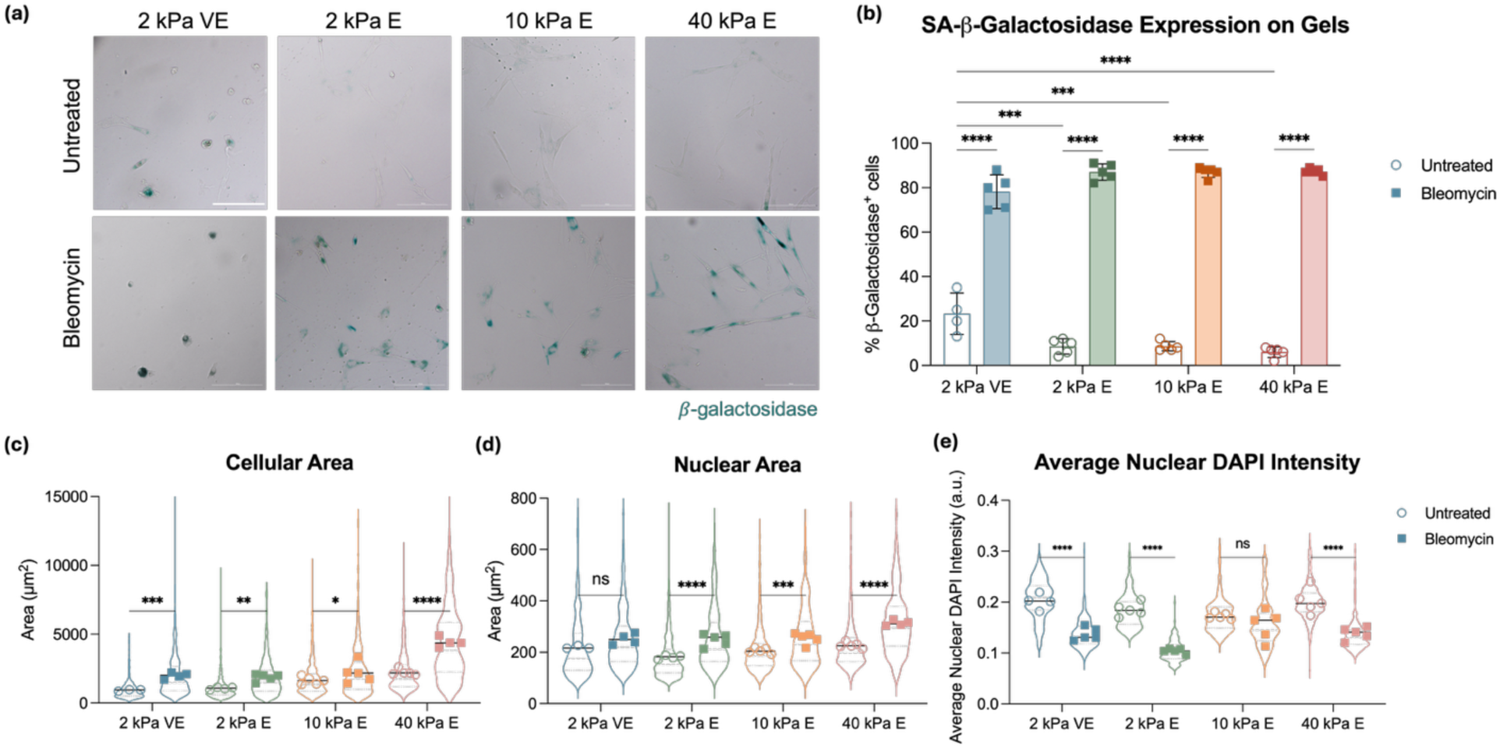
Bleomycin-treated fibroblasts seeded on hydrogels maintain their senescent characteristics. (a) Representative images of fibroblasts either untreated or treated with 20 µg/mL bleomycin prior to seeding onto hydrogels for 24 hours. Scale bar: 200 µm. (b) Cells on all substrates maintained high levels of β-gal staining. Untreated cells seeded on 2 kPa viscoelastic hydrogels also showed a significantly higher level of β-gal activity than those on any other of the hydrogel groups. N = 4-5 hydrogels per group with 327-767 cells per group manually counted. (c) Cell area and (d) nuclear area also significantly increased with bleomycin treatment, apart from the nuclear area of cells on viscoelastic substrates. (e) Average nuclear DAPI intensity also decreased with bleomycin treatment, but there was no statistical difference for cells on 10 kPa hydrogels. Two-way ANOVAs were performed with Tukey’s multiple comparisons tests. N = 3-5 hydrogels per group with 367-1501 cells per group quantified. *p<0.0332, ** p<0.0021, ***p< 0.0002, ****p< 0.0001.

### Viscoelastic substrates increase cellular susceptibility to senescence

Having confirmed that substrate mechanics do not alter an established senescent phenotype, we next evaluated whether substrate mechanics influence fibroblast susceptibility to becoming senescent. Expecting hydrogels to better support senescence induction than tissue culture plastic, we used a 2-day treatment with bleomycin followed by 1 day in normal media, instead of the 3-day treatment used for previous experiments (Fig. 5a). Following this 3-day culture on hydrogels, we fixed and stained for β-gal activity. All hydrogel groups showed significantly higher levels of β-gal expression with bleomycin treatment than the untreated controls, with cells on viscoelastic substrates showing significantly higher β-gal activity than any of the other hydrogel groups and the 40 kPa stiffness group showing significantly lower β-gal activity (Fig. 5b-c). Surprisingly, however, viscoelastic substrates induced senescence to the same degree whether cells were exposed to genotoxic stress or not. Further, the intermediate stiffness group (10 kPa) had non-negligible levels of senescence in the absence of bleomycin treatment.

**Figure 5:**
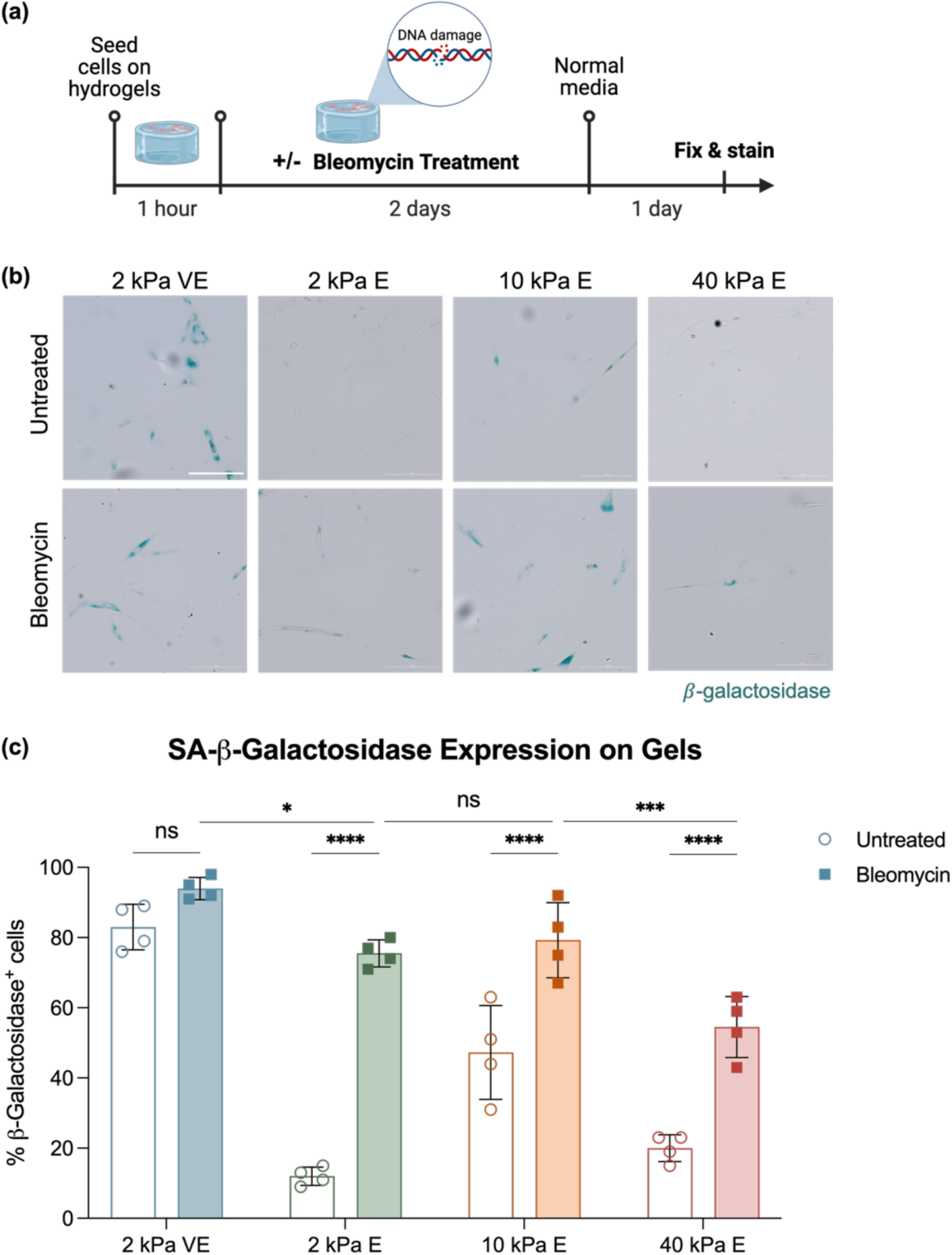
Viscoelastic substrates increase fibroblast susceptibility to bleomycin-induced senescence. (a) Schematic depicting experimental timeline. (b) Representative images depicting β-Gal activity of cells seeded on hydrogels for 3 days. Scale bar: 200 µm. (c) There was no statistical difference in the number of β-Gal^+^ cells between bleomycin-treated and untreated cells on 2 kPa viscoelastic substrates. There was a reduction in the amount of β-gal^+^ cells on all elastic substrates exposed to bleomycin with a significant dip for cells on 40 kPa hydrogels. Untreated cells on 10 kPa hydrogels also showed an increase in β-gal expression relative to the other elastic groups. Two-way ANOVA was performed with Tukey’s multiple comparisons tests. N = 4 hydrogels per group with 488-786 cells per group quantified. *p<0.0332, ***p< 0.0002, ****p< 0.0001.

### Viscoelastic substrates support early markers of senescence following bleomycin treatment

To investigate further this divergence in mechanically-supported senescence induction, we assessed markers of senescence at earlier time points. We seeded fibroblasts on hydrogels and treated with bleomycin for either 12, 24, or 48 hours before fixing and staining for either β-gal or immunostaining for p21 expression and nuclear DAPI intensity (Fig. 6a). At the 12 hour time point, there was significantly higher β-gal expression in cells on 2 kPa viscoelastic substrates compared to 10 kPa elastic hydrogels (Fig. 6b). By 24 hours, cells on 2 kPa viscoelastic substrates had significantly higher β-gal activity than any of the other groups. This trend continued, with the 2 kPa viscoelastic group supporting the highest levels of β-gal activity at 48 hours. While the elastic groups did not show any difference in the amount of β-gal^+^ cells at 12 or 24 hours, by 48 hours each of these groups did show a significant increase in β-gal expression with bleomycin treatment. We also evaluated characteristics of senescent cells including p21 expression and change in nuclear morphology as measured by average nuclear DAPI intensity. Cells on 2 kPa viscoelastic substrates showed significantly depressed nuclear DAPI intensity compared to the other groups at each time point (Fig. 6c). The lower DAPI signal on viscoelastic hydrogels emerged within 12 hours and did not significantly change over 48 hours, suggesting that this could be an early marker of senescence development. Interestingly, the 2 kPa elastic group showed the highest nuclear DAPI signal of the groups at 12 hours, which then decreased at 24 hours to roughly that of the higher stiffness groups. Additionally, cells on 2 kPa viscoelastic substrates showed high levels of p21 expression at each time point (Fig. 6d). This increase in p21 expression on viscoelastic hydrogels was observed at 12 hours and continued to increase at 24 hours. For the elastic hydrogel groups, p21 expression inversely scaled with stiffness. Representative images are shown in Fig. S3.

**Figure 6:**
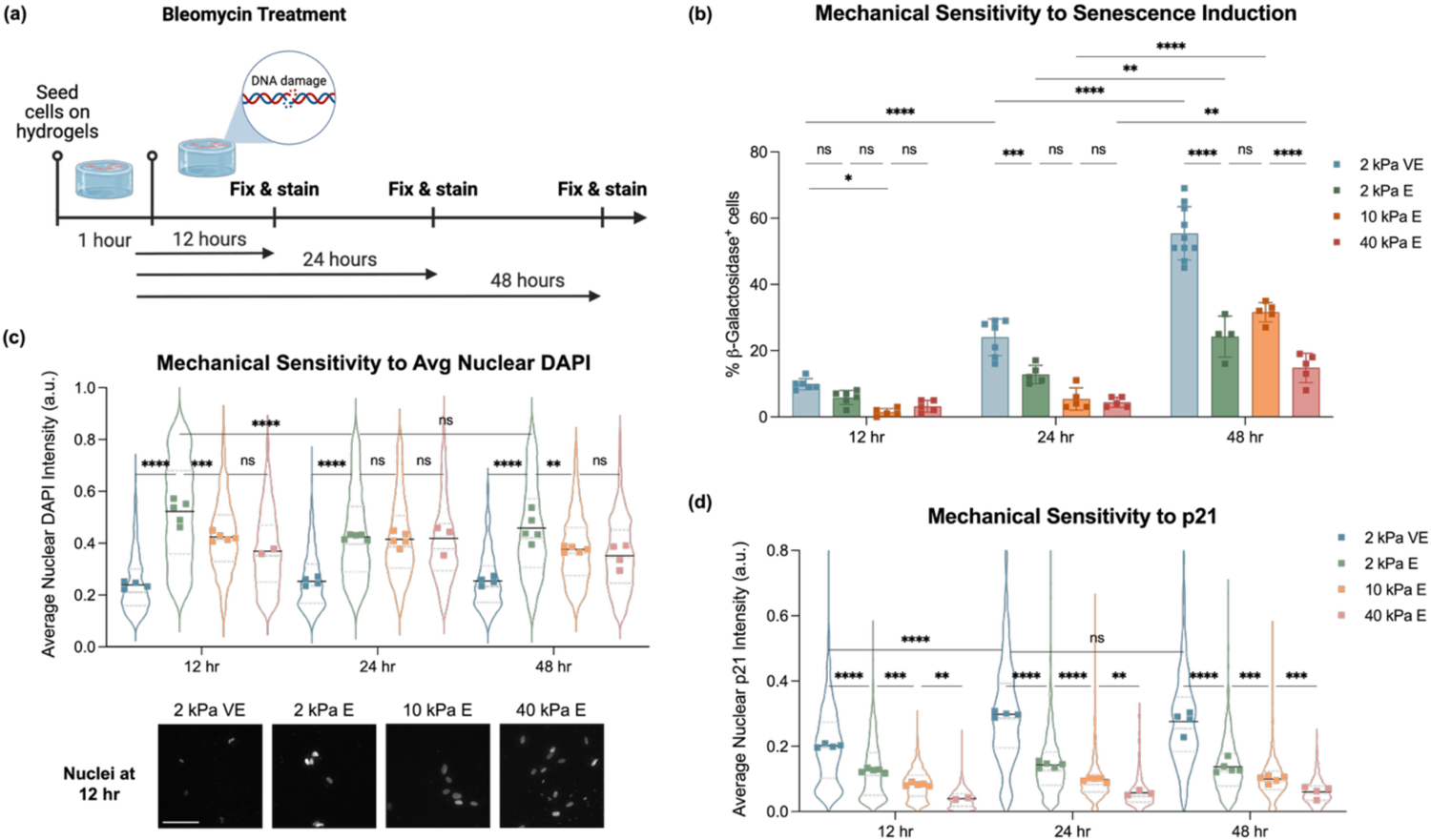
Viscoelastic substrates support early markers of senescence with bleomycin treatment. (a) Schematic illustrating the experimental timeline. (b) Quantification of β-gal expression revealed a significant increase in β-gal^+^ cells on viscoelastic hydrogels. For all groups, β-gal expression increased at 48 hours. N = 4-10 hydrogels per group with a range of 983-2686 cells manually counted per group. (c) Average nuclear DAPI intensity was significantly lower in cells on viscoelastic substrates and did not significantly change with time. Cells on 2 kPa elastic substrates, however, showed high levels of nuclear DAPI that decreased between 12 and 24 hours. Representative nuclei images at 12 hours shown. Scale bar: 100 µm. (d) p21 expression was significantly higher in cells on viscoelastic substrates at 12 hours and continued to increase at 24 hours. In the elastic groups, p21 expression increased with decreasing stiffness. N = 2-5 hydrogels per group with a range of 281-1624 cells per group analyzed. Two-way ANOVAs were performed with Tukey’s multiple comparison tests. *p<0.0332, ** p<0.0021, ***p< 0.0002, ****p< 0.0001.

### Viscoelasticity alone is sufficient to induce cellular senescence

Finally, we assessed early time points and markers of senescence for fibroblasts seeded on hydrogels in the absence of bleomycin-induced genotoxic stress. Just as before, we seeded cells on hydrogels for either 12, 24 or 48 hours before fixing and staining for β-gal or p21 and nuclear DAPI intensity (Fig. 7a). As early as 12 and 24 hours, there was a significant increase in β-gal activity in cells seeded on 2 kPa viscoelastic hydrogels compared to those on 10 kPa or 40 kPa elastic substrates (Fig. 7b). While the other groups did not show a change in the percent of β-gal^+^ cells between time points, at 48 hours the fraction of β-gal^+^ cells on 2 kPa viscoelastic substrates increased significantly. The average DAPI intensity for cells on 2 kPa viscoelastic hydrogels was depressed at 12 hours and did not change significantly over the time points assessed (Fig. 7c). At every time point, cells on 2 kPa viscoelastic substrates showed significantly lower levels of average DAPI than cells on any other hydrogel group. At 48 hours, p21 expression of cells on 2 kPa viscoelastic substrates was significantly higher compared to other experimental groups (Fig. 7d). Representative images are shown in Fig. S4.

**Figure 7:**
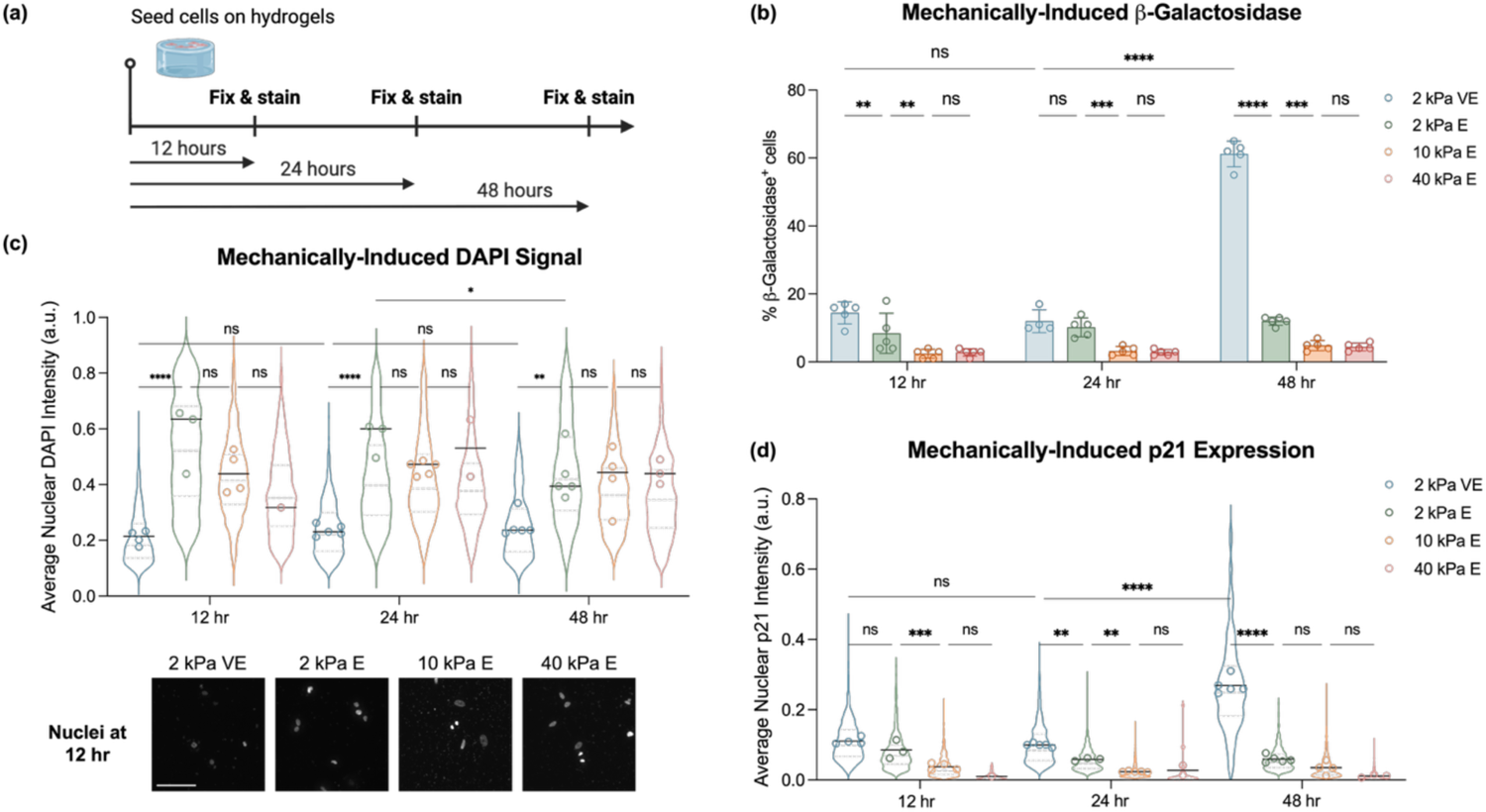
Viscoelasticity alone is sufficient to induce cellular senescence. (a) Schematic representation of experimental timeline. (b) Viscoelastic substrates triggered significantly higher levels of β-gal expression at 48 hours. N = 4-5 hydrogels per group with 1788-2712 cells manually counted per group. (c) Average nuclear DAPI intensity was significantly lower in cells on 2 kPa viscoelastic substrates compared to any other group starting at 12 hours, and did not change from 12 hours to 24 or 48 hours. Representative nuclei images at 12 hours shown. Scale bar: 100 µm. (d) p21 expression was significantly higher in cells on viscoelastic substrates compared to other groups at 48 hours. N = 1-5 hydrogels per group with 28-1429 cells analyzed per group. Two-way ANOVAs were performed with Tukey’s multiple comparison tests. *p<0.0332, ** p<0.0021, ***p< 0.0002, ****p< 0.0001.

## Discussion

This work leveraged IMR-90 primary human lung fibroblasts treated with bleomycin to investigate senescent cell mechanosensitivity. IMR-90 fibroblasts have been thoroughly characterized for their ability to undergo stable senescence, making them a reliable testbed for investigating senescence induction^3,30^. Our evaluation of bleomycin as a method for inducing senescence aligned with what others have seen^15,28,29^, namely that bleomycin treatment induces a robust senescent phenotype through genotoxic stress (Fig. 1). While our work supports previous findings that YAP activity globally declines with senescence^8–10^ (Fig. 2), we further demonstrated that senescent fibroblasts retain their mechanosensing capabilities, showing increased YAP nuclear localization on stiffer hydrogel substrates (Fig. 3). Most notably our results demonstrated that soft viscoelastic hydrogels, with or without genotoxic stress via bleomycin treatment, could potently activate a senescent phenotype in fibroblasts (Figs. 5-7).

These data make clear a unique and potent role of soft viscoelastic substrates in the induction of senescence, independent of genotoxic stress. Given that our viscoelastic hydrogel formulations include a unique polymer modification and crosslinking type, there is the chance that this behavior is triggered by the cell’s response to the guest-host hydrogel chemistry. This is unlikely, however, as cells are not interacting directly with these molecules, and various reports have utilized these chemistries with high cellular viability in 3D culture^26,27,31^, suggesting an even smaller influence on cellular health in 2D culture. We had hypothesized that a decline in YAP activity would support an increase in senescence, in line with what others had reported^8,10^. This alone cannot explain the unique role of viscoelasticity, as we observed equal levels of YAP nuclear localization in stiffness-matched 2 kPa elastic and viscoelastic substrates (Fig. 3c). Thus, the influence of viscoelasticity on senescence induction is most likely due to nuclear changes as suggested by the depression in average nuclear DAPI intensity we observed.

Nuclear wrinkling^32^, weakening of the nuclear envelope^33^, and an overall increase in chromatin accessibility^34,35^ have all been implicated as drivers of senescence. Recent findings demonstrated that soft viscoelastic substrates, in particular, trigger epigenetic remodeling that supports an open chromatin structure^36^. This paper showed that 2 kPa viscoelastic stress-relaxing hydrogels triggered a significant decrease in chromatin compaction compared to elastic 2 kPa hydrogels^36^. Viscoelastic substrates were shown to induce nuclear lamina remodeling by the downregulation of lamin A/C and significant wrinkling in fast-relaxing and slow-relaxing substrates^36^. While the viscoelastic hydrogels used in our work exhibit much faster stress relaxation (Fig. S5) than the viscoelastic substrates from this previous study^36^, the pronounced senescence induction we observed may be driven by nuclear remodeling. However, further studies are needed to elucidate this mechanism more thoroughly.

Elastic substrates appeared to show a non-monotonic relationship between stiffness and senescence induction (Fig. 5c). Again, this does not corroborate the hypothesis that changes in YAP activity drive cellular susceptibility toward senescence, as we know that YAP activity scales with stiffness (Fig. 3c). Thus, nuclear integrity may be similarly playing more of a causative role here than YAP activity. Studies have shown that high levels of tension on the nuclear membrane can lead to rupturing^37,38^, but that high levels of mechanical stimuli can also induce nuclear envelope stiffening^39,40^. Thus, at high stiffnesses, nuclear stiffening may protect against a loss in integrity and senescence, while at low stiffnesses there is no threat to nuclear integrity despite low levels of YAP activity. Further investigation is needed to confirm this direct mechanistic link.

Overall, these results emphasize the importance of viscoelasticity in regulating fibroblast senescence and the need to assess changes in tissue viscoelasticity with age and disease. Given all of our tissues exhibit viscoelastic behavior^17^, it is plausible that the relationship between viscoelasticity and senescence induction is more complex *in vivo* than what is presented in this work. Thus, these results warrant future investigation into the influence of relaxation timescales as well as stiffer viscoelastic substrates on senescence induction, given these mechanical regimes may be more similar to what is found in aged and diseased (e.g., fibrotic^41^) microenvironments.

## Conclusions

This study demonstrates that substrate mechanics – particularly viscoelasticity and stiffness – play a crucial role in the induction of fibroblast senescence. Using a tunable hydrogel platform, we found that soft 2 kPa viscoelastic substrates uniquely support both the onset and amplification of senescent markers, even in the absence of genotoxic stress. Our data reveal that while YAP activity scales with stiffness, its suppression is not solely responsible for initiating senescence, particularly in the context of soft viscoelastic substrates. These findings shed light on the nuanced role that stiffness plays in the induction of senescence and emphasize the underappreciated role of viscoelasticity.

## Methods

### Cell culture

IMR-90 primary lung fibroblasts (ATCC) were grown in Gibco Dulbecco’s Modified Eagle Medium (DMEM) supplemented with 10 v/v% fetal bovine serum (FBS) and 1 v/v% antibiotic-antimycotic (1000 U/mL penicillin, 1000 μg/mL streptomycin, and 0.25 μg/mL amphotericin B) and used between passage 2 and 6 for all experiments. Bleomycin sulfate (ThermoFisher) was used to induce senescence by adding 10 mg/mL stock directly to the cell culture medium to achieve the desired final concentration. Cell culture media was not changed during the duration of the treatment period.

### Senescence-associated β-galactosidase staining

Senescence-associated β-galactosidase was detected using a staining kit following the manufacturer’s instructions (Cell Signaling Technology, MA, #9860). Briefly, cells were fixed for 10 min using a 1X dilution of the provided fixative solution. To prepare the stain, X-Gal (5-bromo-4-chloro-3-indolyl β-D-galactopyranoside) was dissolved at 20 mg/mL in dimethylsulfoxide (DMSO) and combined with the remaining staining components provided at their proper dilutions. The entirety of the staining solution was pH adjusted to 6.0 using either 1 M HCl or 1 M NaOH. A sufficient volume of stain to cover samples thoroughly was added and the plate was sealed with Parafilm and incubated at 37°C overnight. The plate was checked for successful stain development, then the stain was removed and samples were imaged within 3 days. In the case of crystals forming in the plate with an excess of X-Gal, the plate was washed for 10 min with a solution of 50% DMSO and 50% water before proceeding.

### Fluorescent staining

Cells were fixed using 10% neutral-buffered formalin for 15 min and permeabilized with 0.1% Triton X-100 in PBS for 10 min. In the case of immunofluorescent staining, cells were blocked with 3 w/v% bovine serum albumin (BSA) for at least 2 h at room temperature. Primary antibodies were incubated overnight at 4°C while shaking and then washed with PBS. Secondary antibodies and rhodamine phalloidin (Thermo Fisher, 1:400) were incubated at room temperature and protected from light while shaking for at least 2 h, then washed with PBS. Cells were stained with DAPI (Invitrogen, 1:1,000) for 20 minutes before washing with PBS. Antibodies used and their relevant dilution amounts are listed in **Table 1**.

**Table 1:** Antibodies used in this study.

| **Antibody** | **Manufacturer** | **Dilution** |
| --- | --- | --- |
| Anti-Ki67 antibody [B56] (ab279653) | Abcam | 1:200 |
| Anti-p21 antibody [EPR362] | Abcam | 1:200 |
| YAP antibody (63.7) sc-101199 | Santa Cruz Biotech | 1:200 |
| Alexa Fluor 488 goat anti-mouse IgG (H+L) | Thermo Fisher | 1:400 |
| Alexa Fluor 488 goat anti-rabbit IgG (H+L) | Thermo Fisher | 1:400 |
| Alexa Fluor 647 goat anti-mouse IgG (H+L) | Thermo Fisher | 1:400 |
| Alexa Fluor 647 donkey anti-rabbit IgG (H+L) | Thermo Fisher | 1:400 |

### Automatic imaging and analysis

Experiments were imaged using the Cytation C10 confocal imaging system (Agilent, BioTek) at 20X magnification. At least 10 beacons per well were set and captured using autofocus capabilities. For hydrogel experiments, z-stack images were taken and maximum projection images calculated, which were used for subsequent analysis. Cell measurements were taken using a custom CellProfiler pipeline (Broad Institute, Harvard/MIT). Cell nuclei were identified using Otsu adaptive thresholding of the DAPI images, and the propagation of these nuclei were used to identify cell cytoplasm in the rhodamine phalloidin images. Measurements of cell shape, texture, and stain intensity were then taken on these objects for the relevant images and used to create quantitative outputs. For quantification of the percent of β-gal^+^ cells, two independent researchers manually counted the number of cells with positive staining using color brightfield images and manually counted the number of corresponding cells using DAPI-stained fluorescent images. The average of the two counts were then plotted for each group.

### RT-qPCR

RNA isolation was performed using an RNeasy Mini Kit (Qiagen, 74904) according to the manufacturer’s instructions on cells that had been removed from their culture surface. RNA quantification and quality assessment was performed using a Nanodrop 2000 spectrophotometer.

We discarded samples with 260/280 nm ratio of less than 1.7 or a 260/230 nm ratio of less than 1. We then synthesized cDNA using a QuantiTech Reverse Transcriptase kit with genomic DNA removal (Qiagen), following instructions provided by the manufacturer and using an input of 450 ng of RNA. We then proceeded to qPCR using iQ™ SYBR® Green Supermix (BioRad, 1708880) and a BioRad iCycler. The primers used are listed in **Table 2**. All qPCR reactions were run in duplicate with the values averaged and then normalized to *GAPDH* expression levels.

**Table 2:**
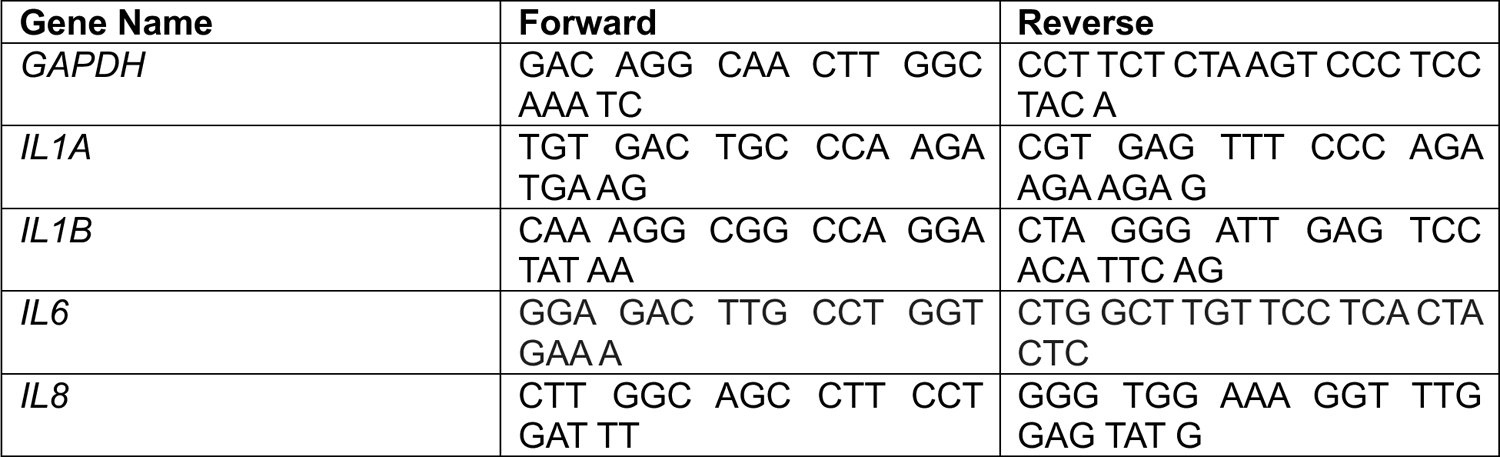
Primers used in this study.

### NorHA synthesis

As previously reported^42^, hyaluronic acid was functionalized with norbornene groups via activation of the carboxylic acid using 4-(4,6-dimethoxy[1.3.5]triazin-2-yl)-4-methylmorpholinium chloride (DMTMM), and subsequent amine substitution of 5-norbornene-2-methylamine. Specifically, sodium hyaluronate (NaHA) (Lifecore, 82 kDa) and 2-(N-Morpholino)ethanesulfonic acid (MES) were dissolved in deionized (DI) water at a 1:1.15 ratio and adjusted to a pH of 5.5 using 10 M NaOH. Then, DMTMM was dissolved into the solution at a ratio of 1.24 g to 1 g NaHA. 5-norbornene-2-methlyamine was added dropwise and allowed to stir overnight. The solution was then precipitated using cold ethanol. The precipitate was collected via vacuum filtration and dissolved in 2 M brine for dialysis against water for 1 day and 1 M brine for an additional day. The dialysis process was repeated three times before the product was frozen and lyophilized for long-term storage at −20°C. The final product was analyzed through ^1^H NMR and the degree of norbornene modification was determined to be 23% (Fig. S6).

### CDHA synthesis

β-cyclodextrin-modified hyaluronic acid (CDHA) was synthesized as previously described^25,43^. Briefly, β-cyclodextrin hexamethylene diamine (CDHDA) was made by adding dropwise p-Toluenesulfonyl chloride (TosCl) dissolved in acetonitrile to a suspension of CD in ultra-pure water (1.25:1 molar ratio of TosCl to CD) over ice. After stirring for 2 h, cold NaOH was added dropwise (3.1:1 molar ratio of NaOH to CD) to the solution, then the reaction was left for an additional 30 min off ice. Ammonium chloride was added until a pH of 8.5 was reached, then the solution was cooled on ice and centrifuged to recover the precipitate. Cold water was used to decant and reprecipitate 3 times, then the product was washed 3 times in cold acetone and once with cold diethyl ether and left to dry overnight. The CD-Tos product was charged with HDA (4 g/g CD-Tos) and dimethylformamide (DMF) (5 mL/g CD-Tos) and left to react for 12 h at 80°C under nitrogen. The final CDHDA product was precipitated with cold acetone, washed with cold diethyl ether, and vacuum dried. The degree of modification was determined to be 73% by ^1^H NMR^27^. To achieve the final CDHA polymer, CDHDA was reacted with HA-TBA and BOP in anhydrous DMSO at 25°C for 3 h. This reaction was quenched with cold water, dialyzed for 5 days, filtered, and dialyzed for 5 more days. The purified solution was frozen and lyophilized. The degree of modification of HA was determined by ^1^H NMR to be 25%^27^ (Fig. S7).

### HA hydrogel fabrication

As has been previously described, hydrogels were formed through ultraviolet (UV)-light mediated thiol-ene addition^25,27,44^. Elastic hydrogels were formed using dithiothreitol (DTT) crosslinks and varying the weight percent of NorHA. Stiffnesses of 2 kPa (2 wt%), 10 kPa (3 wt%) and 40 kPa (6 wt%) were achieved by using a thiol:norbornene ratio of 0.2, 0.4, and 0.4 respectively. Viscoelasticity was introduced by incorporating physical interactions between CDHA and a thiolated adamantane peptide (GCKKK-Adamantane, Genscript)^25^. Viscoelastic hydrogels with tan delta (loss to storage modulus ratio) ∼ 0.15 (4 wt% CDHA-NorHA) were crosslinked using DTT at a thiol:norbornene ratio of 0.1 and a CDHA-adamantane peptide mixture at a CD:adamantane ratio of 1.3:1. Cell adhesion was enabled through the incorporation of 1 mM RGD peptide (GCGYG<u>RGD</u>SPG, Genscript) in all hydrogel groups via thiol-ene addition of the thiol on the peptide’s cysteine to a free norbornene. UV-light mediated polymerization was enabled by the incorporation of 1 mM lithium acylphosphinate (LAP) photoinitiator into each hydrogel precursor solution.

For hydrogel experiments conducted in 96-well plates, these plates were fabricated as previously described^27^. Briefly, a glass piece with the dimensions of a 96 well plate was treated with 95% (3-mercaptopropyl)trimethoxysilane to conjugate free thiols onto the surface, baked at 100°C for 1 h, and sequentially washed with dichloromethane (DCM), 70% ethanol, and DI water. A silicone spacer sheet cut to the dimensions of a 96-well plate (Grace BioLabs) was then adhered to this glass piece and 18 µL of the hydrogel precursor solutions were added to each well. These precursor solutions were then flattened using another glass piece treated with the hydrophobic coating SigmaCote before placing in a UV box (VWR) for photopolymerization (365 nm, 5 mW/cm^2^) for 2 min. The top glass piece and silicone spacer sheet were then removed and a bottomless 96 well plate with double-sided adhesive was applied to the functionalized glass piece. Hydrogels within the wells were then swollen with PBS at 37°C and sterilized using 2 h of germicidal UV before utilizing for experiments.

For hydrogel experiments conducted in 6-well plates, a similar approach was taken using square 22 mm x 22 mm coverslips. Individual coverslips were treated with 95% (3-mercaptopropyl)trimethoxysilane and baked at 100°C for 1 h before washing with DCM, 70% ethanol, and DI water. Hydrogel precursor solutions of 50 µL were pipetted onto the glass surface and another 18 mm x 18 mm coverslip was used to flatten solution before UV-curing (5 mW/cm^2^, 2 min). Once curing was complete and a thin hydrogel film had formed, the top cover glass was removed using a razor blade and the hydrogel-bound cover glass was moved into a 6-well plate (Thermo Fisher) for sterilization.

### Mechanical characterization

Hydrogel mechanics were assessed via nanoindentation on hydrogels swollen in PBS overnight at 37°C using Optics 11 Life Piuma nanoindenter. A 50 µm borosilicate glass probe attached to a cantilever with a spring constant of 0.5 N/m was used for all tests. Indentations were made with a depth of 5 µm after the surface was recognized increasing depth at a constant rate for 2 sec. The Hertzian contact model was applied and a Poisson’s ratio of 0.5 was assumed. For each hydrogel, a matrix of 4 points was tested, with a distance of 200 µm between each point, and the average of the four points was taken to represent each hydrogel. Average hydrogel mechanics are plotted in Fig. S5. Additionally, stress relaxation tests were run to confirm viscoelastic character in hydrogels containing guest-host crosslinks. These tests were run with the same indentation parameters as before, but held at a constant depth for 30 sec before removing. Stress was normalized to the maximum stress and plotted over time (Fig. S5)

### Statistical analysis

Data plotting and statistical analysis was performed using GraphPad Prism (v. 10.3.1). For each graph, points plotted represent data from a unique well or hydrogel in which all individual cellular measurements taken from that well were averaged. Overlaid violin plots represent all individual cellular measurements for all replicates in a given group. For comparisons between bleomycin-treated and untreated groups, t-tests were used and statistically significant differences are indicated by *, **, ***, **** corresponding to p < 0.0332, 0.0021, 0.0002, or 0.0001 respectively. Additional details on sample size or alternative statistical tests are included in figure captions.

## Resource Availability

### Lead Contact

Further information and requests for resources and reagents should be directed to and will be fulfilled by the lead contact, Steven Caliari.

### Materials Availability

Hydrogel materials (NorHA, CDHA, etc.) in this study will be made available on request, but we may require a payment and/or a completed materials transfer agreement if there is potential for commercial application.

### Data and Code Availability

All data needed for the conclusions in the paper are present in the paper and the supplemental information. Additional data for the paper are available upon reasonable request from the lead contact.

## Acknowledgments

The authors would like to thank James Gentry for providing the norbornene-modified hyaluronic acid utilized in this work and for helpful discussion on experimental design; Leilani Astrab for assistance with β-cyclodextrin-modified hyaluronic acid synthesis, plotting of ^1^H NMR spectra, and invaluable feedback on experimental and figure design; and Manasi Krishnakumar for running ^1^H NMR on samples. This work was supported by the NIH (R35GM138187) and NSF (CAREER DMR/BMAT 2046592, GRFP to M.L.S.). The content is solely the responsibility of the authors and does not necessarily represent the official views of the National Institutes of Health.

## Author Contributions

Conceptualization, M.L.S and S.R.C.; Methodology, M.L.S and S.R.C.; Investigation, M.L.S., T.B., E.Y., and A.O.; Writing – Original Draft, M.L.S.; Writing – Review & Editing, M.L.S and S.R.C.; Funding Acquisition, M.L.S. and S.R.C.

## References

1. Childs, B.G., Durik, M., Baker, D.J., and van Deursen, J.M. (2015). Cellular senescence in aging and age-related disease: from mechanisms to therapy. Nat. Med. 21, 1424–1435. 10.1038/nm.4000.

2. Muñoz-Espín, D., and Serrano, M. (2014). Cellular senescence: from physiology to pathology. Nat. Rev. Mol. Cell Biol. 15, 482–496. 10.1038/nrm3823.

3. Schafer, M.J., White, T.A., Iijima, K., Haak, A.J., Ligresti, G., Atkinson, E.J., Oberg, A.L., Birch, J., Salmonowicz, H., Zhu, Y., et al. (2017). Cellular senescence mediates fibrotic pulmonary disease. Nat. Commun. 8, 14532. 10.1038/ncomms14532.

4. Álvarez, D., Cárdenes, N., Sellarés, J., Bueno, M., Corey, C., Hanumanthu, V.S., Peng, Y., D’Cunha, H., Sembrat, J., Nouraie, M., et al. (2017). IPF lung fibroblasts have a senescent phenotype. Am. J. Physiol.-Lung Cell. Mol. Physiol. 313, L1164–L1173. 10.1152/ajplung.00220.2017.

5. Calhoun, C., Shivshankar, P., Saker, M., Sloane, L.B., Livi, C.B., Sharp, Z.D., Orihuela, C.J., Adnot, S., White, E.S., Richardson, A., et al. (2016). Senescent Cells Contribute to the Physiological Remodeling of Aged Lungs. J. Gerontol. Ser. A 71, 153–160. 10.1093/gerona/glu241.

6. Ulldemolins, A., Narciso, M., Sanz-Fraile, H., Otero, J., Farré, R., Gavara, N., and Almendros, I. (2024). Effects of aging on the biomechanical properties of the lung extracellular matrix: dependence on tissular stretch. Front. Cell Dev. Biol. 12. 10.3389/fcell.2024.1381470.

7. Birch, H.L. (2018). Extracellular Matrix and Ageing. In Biochemistry and Cell Biology of Ageing: Part I Biomedical Science, J. R. Harris and V. I. Korolchuk, eds. (Springer), pp. 169–190. 10.1007/978-981-13-2835-0_7.

8. Sladitschek-Martens, H.L., Guarnieri, A., Brumana, G., Zanconato, F., Battilana, G., Xiccato, R.L., Panciera, T., Forcato, M., Bicciato, S., Guzzardo, V., et al. (2022). YAP/TAZ activity in stromal cells prevents ageing by controlling cGAS–STING. Nature, 1–9. 10.1038/s41586-022-04924-6.

9. Joung, J., Heo, Y., Kim, Y., Kim, J., Choi, H., Jeon, T., Jang, Y., Kim, E.-J., Lee, S.H., Suh, J.M., et al. (2025). Cell enlargement modulated by GATA4 and YAP instructs the senescence-associated secretory phenotype. Nat. Commun. 16, 1696. 10.1038/s41467-025-56929-0.

10. Xie, Q., Chen, J., Feng, H., Peng, S., Adams, U., Bai, Y., Huang, L., Li, J., Huang, J., Meng, S., et al. (2013). YAP/TEAD–Mediated Transcription Controls Cellular Senescence. Cancer Res. 73, 3615–3624. 10.1158/0008-5472.CAN-12-3793.

11. Yao, X., Li, H., Chen, L., and Tan, L.P. (2022). UV-induced senescence of human dermal fibroblasts restrained by low-stiffness matrix by inhibiting NF-*κ*B activation. Eng. Regen. 3, 365–373. 10.1016/j.engreg.2022.08.002.

12. Blokland, K.E.C., Nizamoglu, M., Habibie, H., Borghuis, T., Schuliga, M., Melgert, B.N., Knight, D.A., Brandsma, C.-A., Pouwels, S.D., and Burgess, J.K. (2022). Substrate stiffness engineered to replicate disease conditions influence senescence and fibrotic responses in primary lung fibroblasts. Front. Pharmacol. 13. 10.3389/fphar.2022.989169.

13. Du, H., Rose, J.P., Bons, J., Guo, L., Valentino, T.R., Wu, F., Burton, J.B., Basisty, N., Manwaring-Mueller, M., Makhijani, P., et al. (2025). Substrate stiffness dictates unique doxorubicin-induced senescence-associated secretory phenotypes and transcriptomic signatures in human pulmonary fibroblasts. GeroScience. 10.1007/s11357-025-01507-x.

14. He, H., Zeng, B., Wu, X., Hou, J., Wang, Y., Wang, Y., Lin, Y., Wu, P., Zheng, C., Yin, H., et al. (2023). Higher matrix stiffness promotes VSMC senescence by affecting mitochondria–ER contact sites and mitochondria/ER dysfunction. FASEB J. 37, e23318. 10.1096/fj.202301198RR.

15. Starich, B., Yang, F., Tanrioven, D., Kung, H.-C., Baek, J., Nair, P.R., Kamat, P., Macaluso, N., Eoh, J., Han, K.S., et al. (2024). Substrate stiffness modulates the emergence and magnitude of senescence phenotypes in dermal fibroblasts. Preprint at bioRxiv, 10.1101/2024.02.06.579151.

16. Felisbino, M.B., Rubino, M., Travers, J.G., Schuetze, K.B., Lemieux, M.E., Anseth, K.S., Aguado, B.A., and McKinsey, T.A. (2024). Substrate stiffness modulates cardiac fibroblast activation, senescence, and proinflammatory secretory phenotype. Am. J. Physiol.-Heart Circ. Physiol. 326, H61–H73. 10.1152/ajpheart.00483.2023.

17. Chaudhuri, O., Cooper-White, J., Janmey, P.A., Mooney, D.J., and Shenoy, V.B. (2020). Effects of extracellular matrix viscoelasticity on cellular behaviour. Nature 584, 535–546. 10.1038/s41586-020-2612-2.

18. Charrier, E.E., Pogoda, K., Wells, R.G., and Janmey, P.A. (2018). Control of cell morphology and differentiation by substrates with independently tunable elasticity and viscous dissipation. Nat. Commun. 9, 449. 10.1038/s41467-018-02906-9.

19. Elosegui-Artola, A. (2021). The extracellular matrix viscoelasticity as a regulator of cell and tissue dynamics. Curr. Opin. Cell Biol. 72, 10–18. 10.1016/j.ceb.2021.04.002.

20. Huerta-López, C., Clemente-Manteca, A., Velázquez-Carreras, D., Espinosa, F.M., Sanchez, J.G., Martínez-del-Pozo, Á., García-García, M., Martín-Colomo, S., Rodríguez-Blanco, A., Esteban-González, R., et al. (2024). Cell response to extracellular matrix viscous energy dissipation outweighs high-rigidity sensing. Sci. Adv. 10, eadf9758. 10.1126/sciadv.adf9758.

21. Wang, X., Song, L., Zhao, J., Xiong, Y., Jin, R., and He, J. (2025). Matrix viscoelasticity drives cell cluster formation to counteract cellular senescence. J. Mater. Chem. B. 10.1039/D5TB00174A.

22. Aragona, M., Panciera, T., Manfrin, A., Giulitti, S., Michielin, F., Elvassore, N., Dupont, S., and Piccolo, S. (2013). A Mechanical Checkpoint Controls Multicellular Growth through YAP/TAZ Regulation by Actin-Processing Factors. Cell 154, 1047–1059. 10.1016/j.cell.2013.07.042.

23. González-Gualda, E., Baker, A.G., Fruk, L., and Muñoz-Espín, D. (2021). A guide to assessing cellular senescence in vitro and in vivo. FEBS J. 288, 56–80. 10.1111/febs.15570.

24. Mascetti, G., Carrara, S., and Vergani, L. (2001). Relationship between chromatin compactness and dye uptake for in situ chromatin stained with DAPI. Cytometry 44, 113–119. 10.1002/1097-0320(20010601)44:2<113::aid-cyto1089>3.0.co;2-a.

25. Hui, E., Gimeno, K.I., Guan, G., and Caliari, S.R. (2019). Spatiotemporal Control of Viscoelasticity in Phototunable Hyaluronic Acid Hydrogels. Biomacromolecules 20, 4126– 4134. 10.1021/acs.biomac.9b00965.

26. Miller, B., Hansrisuk, A., Highley, C.B., and Caliari, S.R. (2021). Guest–Host Supramolecular Assembly of Injectable Hydrogel Nanofibers for Cell Encapsulation. ACS Biomater. Sci. Eng. 7, 4164–4174. 10.1021/acsbiomaterials.1c00275.

27. Skelton, M.L., Gentry, J.L., Astrab, L.R., Goedert, J.A., Earl, E.B., Pham, E.L., Bhat, T., and Caliari, S.R. (2024). Modular Multiwell Viscoelastic Hydrogel Platform for Two- and Three-Dimensional Cell Culture Applications. ACS Biomater. Sci. Eng. 10, 3280–3292. 10.1021/acsbiomaterials.4c00312.

28. Chen, F., Zhao, W., Du, C., Chen, Z., Du, J., and Zhou, M. (2024). Bleomycin induces senescence and repression of DNA repair via downregulation of Rad51. Mol. Med. Camb. Mass 30, 54. 10.1186/s10020-024-00821-y.

29. Savić, R., Yang, J., Koplev, S., An, M.C., Patel, P.L., O’Brien, R.N., Dubose, B.N., Dodatko, T., Rogatsky, E., Sukhavasi, K., et al. (2023). Integration of transcriptomes of senescent cell models with multi-tissue patient samples reveals reduced COL6A3 as an inducer of senescence. Cell Rep. 42, 113371. 10.1016/j.celrep.2023.113371.

30. Sherwood, S.W., Rush, D., Ellsworth, J.L., and Schimke, R.T. (1988). Defining cellular senescence in IMR-90 cells: a flow cytometric analysis. Proc. Natl. Acad. Sci. 85, 9086–9090. 10.1073/pnas.85.23.9086.

31. Hong, K.H., and Song, S.-C. (2019). 3D hydrogel stem cell niche controlled by host-guest interaction affects stem cell fate and survival rate. Biomaterials 218, 119338. 10.1016/j.biomaterials.2019.119338.

32. Pathak, R.U., Soujanya, M., and Mishra, R.K. (2021). Deterioration of nuclear morphology and architecture: A hallmark of senescence and aging. Ageing Res. Rev. 67, 101264. 10.1016/j.arr.2021.101264.

33. Martins, F., Sousa, J., Pereira, C.D., da Cruz e Silva, O.A.B., and Rebelo, S. (2020). Nuclear envelope dysfunction and its contribution to the aging process. Aging Cell 19, e13143. 10.1111/acel.13143.

34. Criscione, S.W., Teo, Y.V., and Neretti, N. (2016). The chromatin landscape of cellular senescence. Trends Genet. TIG 32, 751–761. 10.1016/j.tig.2016.09.005.

35. Lopes-Paciencia, S., and Ferbeyre, G. (2025). Increased chromatin accessibility underpins senescence. FEBS J. 10.1111/febs.70136.

36. Wu, Y., Song, Y., Soto, J., Hoffman, T., Lin, X., Zhang, A., Chen, S., Massad, R.N., Han, X., Qi, D., et al. (2025). Viscoelastic extracellular matrix enhances epigenetic remodeling and cellular plasticity. Nat. Commun. 16, 4054. 10.1038/s41467-025-59190-7.

37. Zhang, Q., Tamashunas, A.C., Agrawal, A., Torbati, M., Katiyar, A., Dickinson, R.B., Lammerding, J., and Lele, T.P. (2019). Local, transient tensile stress on the nuclear membrane causes membrane rupture. Mol. Biol. Cell 30, 899–906. 10.1091/mbc.E18-09-0604.

38. Baumann, K. (2016). Nuclear envelope ruptures as cells squeeze through tight spaces. Nat. Rev. Mol. Cell Biol. 17, 263–263. 10.1038/nrm.2016.47.

39. Tang, W., Chen, X., Wang, X., Zhu, M., Shan, G., Wang, T., Dou, W., Wang, J., Law, J., Gong, Z., et al. (2023). Indentation induces instantaneous nuclear stiffening and unfolding of nuclear envelope wrinkles. Proc. Natl. Acad. Sci. 120, e2307356120. 10.1073/pnas.2307356120.

40. Swift, J., Ivanovska, I.L., Buxboim, A., Harada, T., Dingal, P.C.D.P., Pinter, J., Pajerowski, J.D., Spinler, K.R., Shin, J.-W., Tewari, M., et al. (2013). Nuclear Lamin-A Scales with Tissue Stiffness and Enhances Matrix-Directed Differentiation. Science 341, 1240104. 10.1126/science.1240104.

41. Long, Y., Niu, Y., Liang, K., and Du, Y. (2022). Mechanical communication in fibrosis progression. Trends Cell Biol. 32, 70–90. 10.1016/j.tcb.2021.10.002.

42. Darling, N.J., Xi, W., Sideris, E., Anderson, A.R., Pong, C., Carmichael, S.T., and Segura, T. (2020). Click by Click Microporous Annealed Particle (MAP) Scaffolds. Adv. Healthc. Mater. 9, 1901391. 10.1002/adhm.201901391.

43. Rodell, C.B., Kaminski, A.L., and Burdick, J.A. (2013). Rational Design of Network Properties in Guest–Host Assembled and Shear-Thinning Hyaluronic Acid Hydrogels. Biomacromolecules 14, 4125–4134. 10.1021/bm401280z.

44. Hui, E., Moretti, L., Barker, T.H., and Caliari, S.R. (2021). The Combined Influence of Viscoelastic and Adhesive Cues on Fibroblast Spreading and Focal Adhesion Organization. Cell. Mol. Bioeng. 14, 427–440. 10.1007/s12195-021-00672-1.

