## Supplemental Information for "Substrate stiffness and viscoelasticity influence fibroblast senescence"

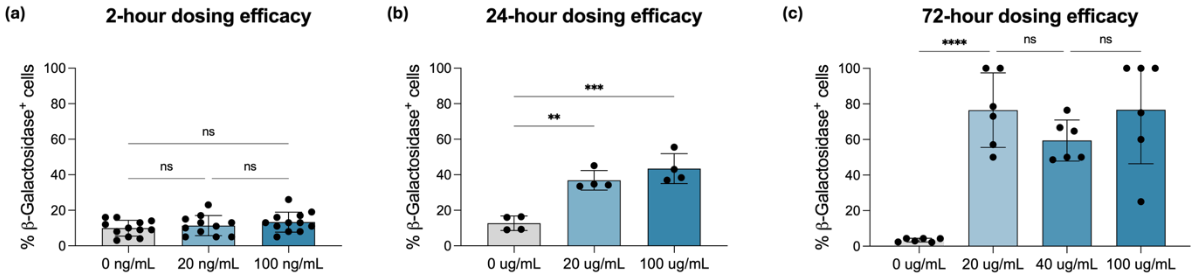


**Supplemental Figure 1: Timing of bleomycin dosing affects senescence induction.** Fibroblasts were treated with varying doses of bleomycin for either 2, 24, or 72 hours before fixing and staining for β-galactosidase (β-gal) expression. (a) Cells treated for 2 hours did not show higher levels of β-gal expression with increasing dose. (b) At 24 hours of treatment, there is a dose-dependent senescence response to bleomycin treatment. (c) At 72 hours of bleomycin treatment, all doses result in high levels of β-gal expression. One-way ANOVA was performed with Tukey's post-hoc analysis. **p< 0.0021, ***p<0.0002.

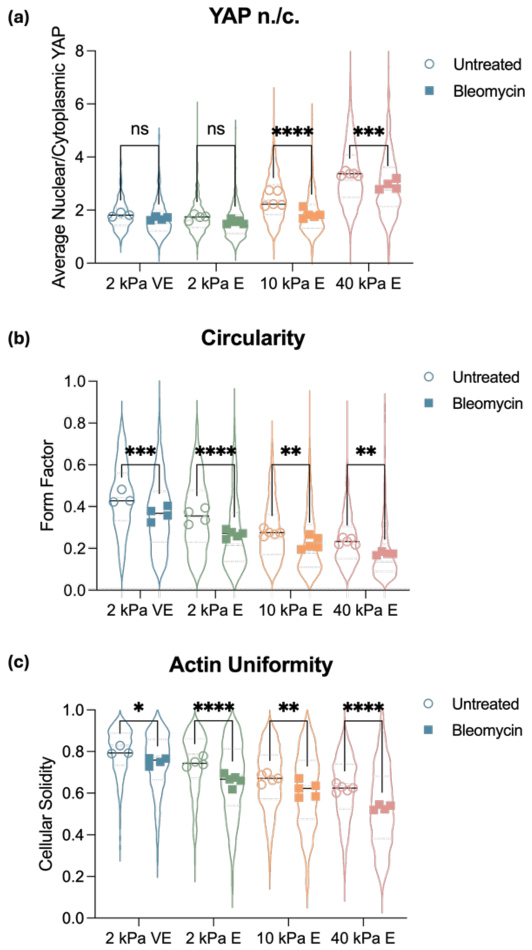


**Supplemental Figure 2: Bleomycin treatment induces differences in YAP activity and cellular morphology on hydrogels.** (a) Bleomycin-treated fibroblasts on 10 and 40 kPa hydrogels showed a significant decrease in YAP nuclear localization relative to the untreated controls. There was a shift downward in YAP activity for cells on 2 kPa substrates, but these differences were not statistically significant. Additionally, bleomycin-treated cells on all substrates showed a significant decrease in (b) circularity and (c) actin uniformity (indicating increased actin fiber organization). Two-way ANOVA tests were performed with Tukey's multiple comparison test. N = 3-5 hydrogels per experimental group with 40-396 cells per individual hydrogel quantified. *p<0.0332, ***p<0.0002, ****p<0.0001.

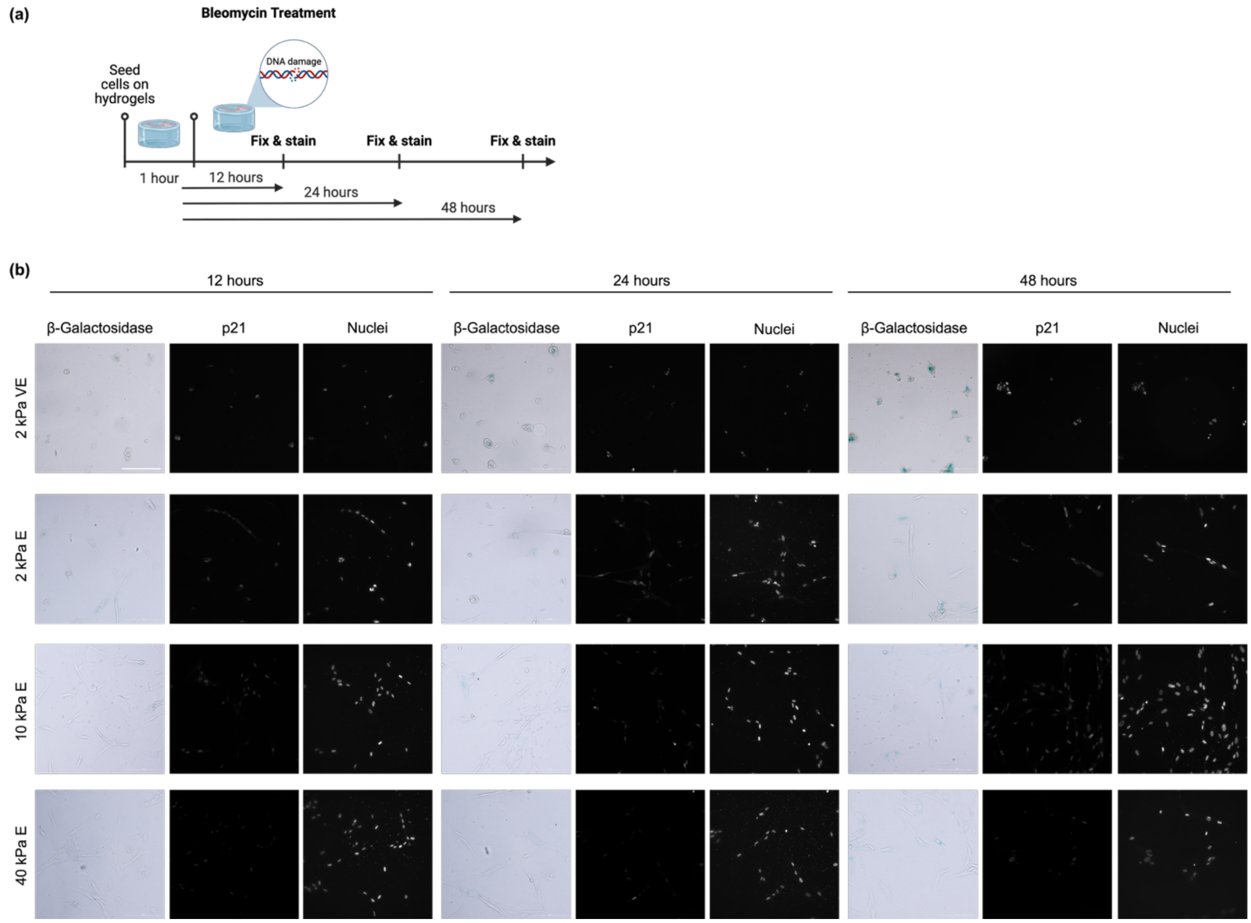


**Supplemental Figure 3: Representative images from data in Figure 6.** (a) Schematic depicting experimental timeline, and (b) representative images of fibroblast β-gal and p21 expression for each time point and hydrogel group in Figure 6. Scale bar: 200 μm.

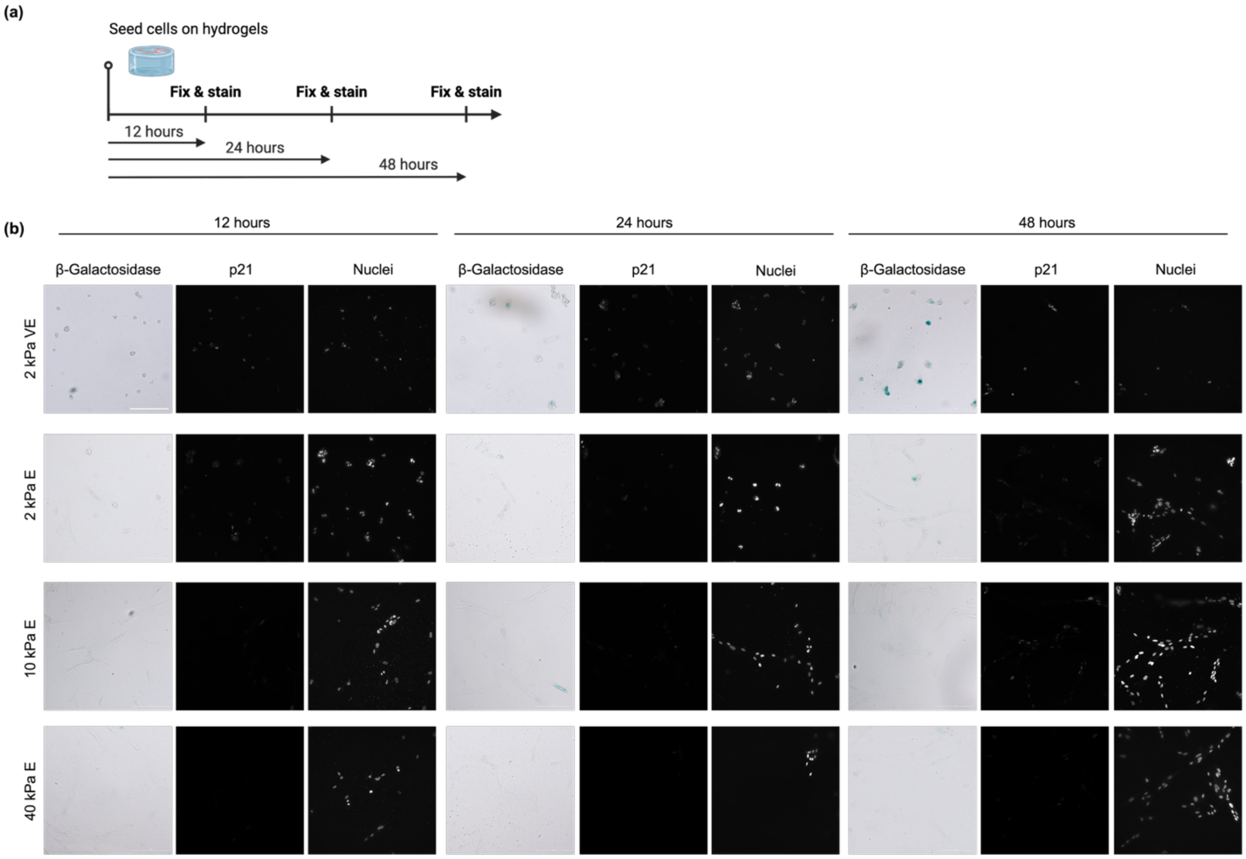


**Supplemental Figure 4: Representative images from data in Figure 7.** (a) Schematic depicting experimental timeline, and (b) representative images of fibroblast β-gal and p21 expression for each time point and hydrogel group in Figure 7. Scale bar: 200 μm.

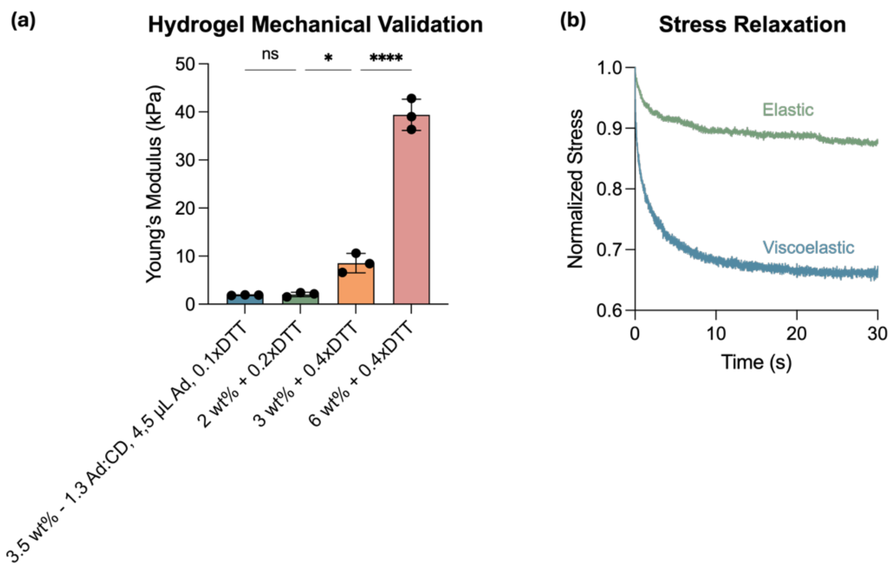


**Supplemental Figure 5: Hydrogel mechanical analysis.** (a) Hydrogel stiffness was tested using nanoindentation and formulations were tuned to achieve a range of stiffnesses. N = 3 hydrogels assessed per group. One-way ANOVA test was performed with Tukey's post-hoc analysis. *$p<0.0332$, ****$p<0.0001$. (b) Stress relaxation tests were run to verify viscoelastic behavior of guest-host hydrogels. Viscoelastic hydrogels show a greater extent of stress relaxation than elastic hydrogels.

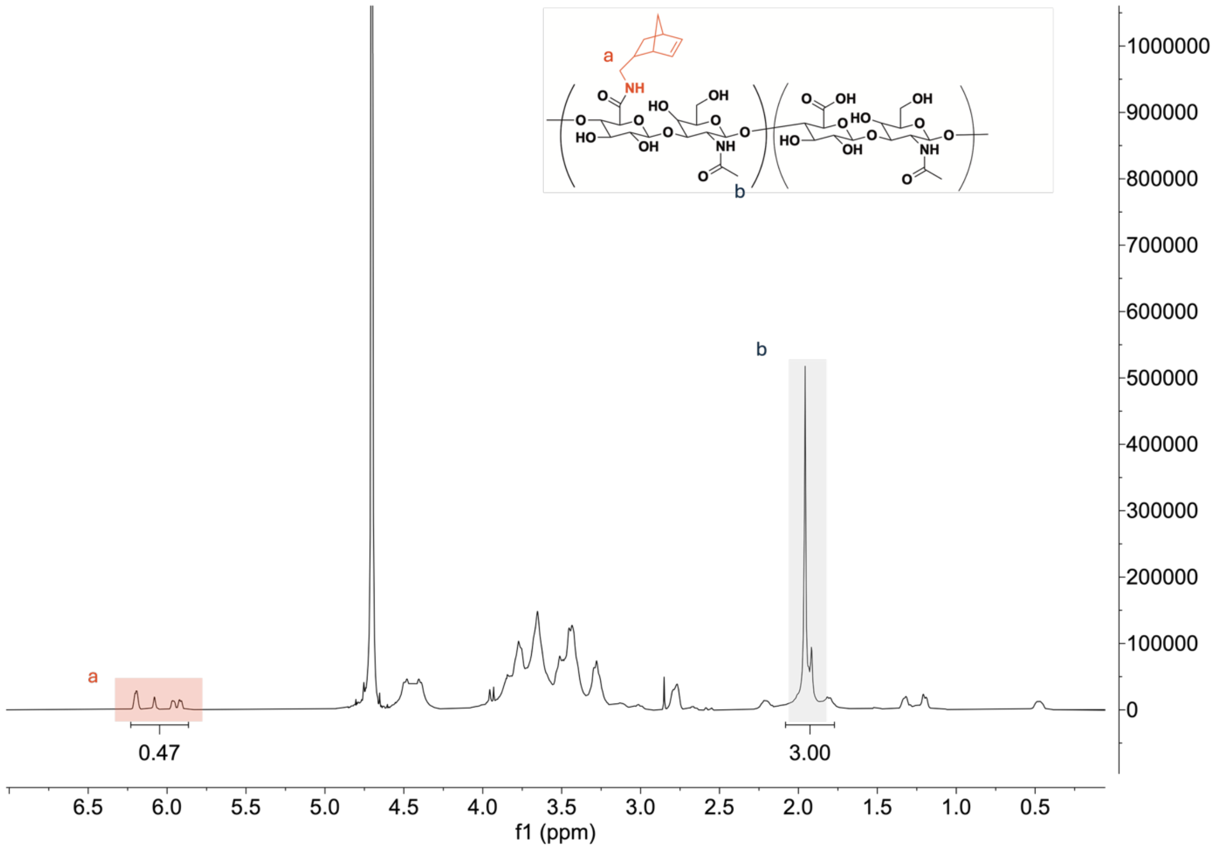


**Supplemental Figure 6: [1]H NMR spectrum of norbornene-modified hyaluronic acid (NorHA).** The degree of modification of HA was determined to be 23.5% as indicated by the integration of the peaks ranging from δ = 5.75 – 6.2 (a) that represent the two hydrogens on either end of the double bond of the norbornene molecule. This was normalized to the three hydrogens on the *N*-acetyl group present on each HA repeat unit (b).

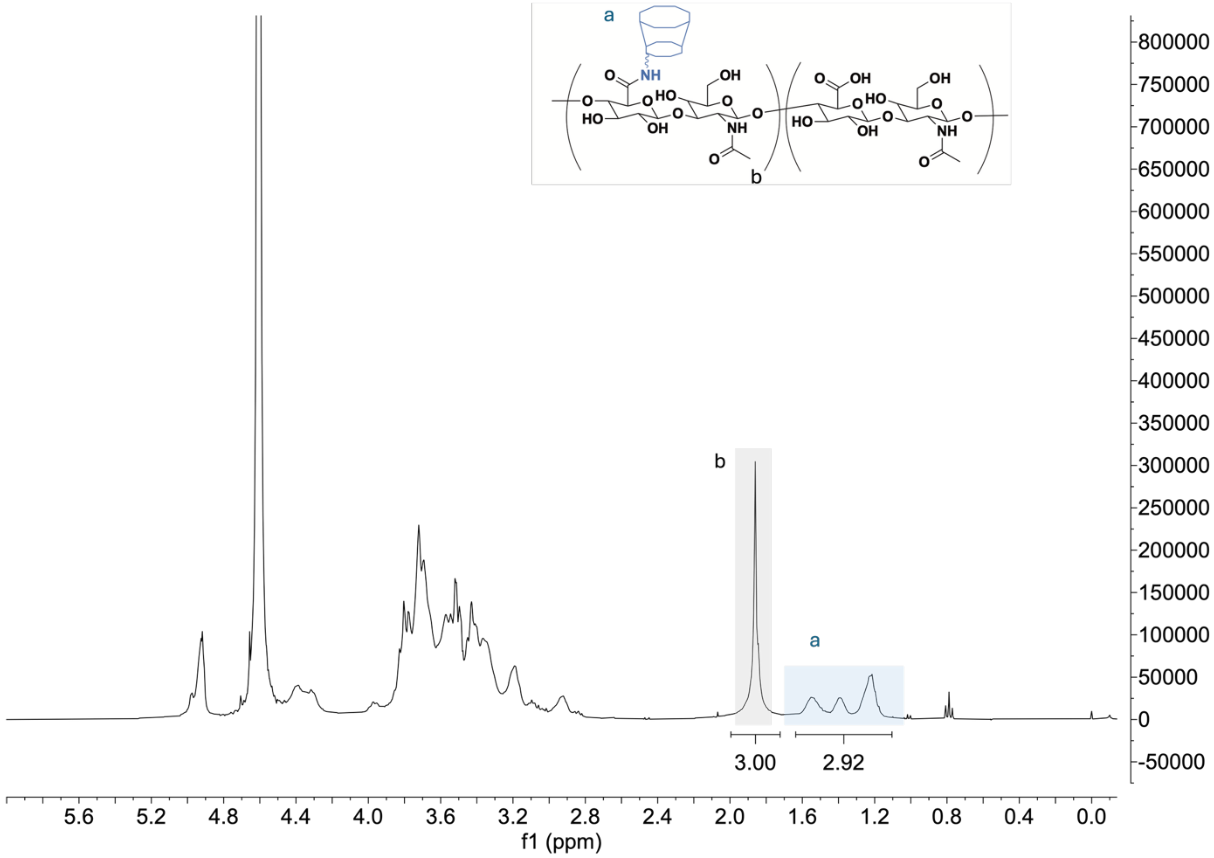


**Supplemental Figure 7: [1]H NMR spectrum of β-cyclodextrin-modified hyaluronic acid (CDHA).** Modification of HA with pendant β-cyclodextrins (25.2%) is determined by integration of hexane linker hydrogens (12H, shaded blue) relative to the *N*-acetyl group of HA (3H, shaded grey).
